# Effects of Chronic Cannabis Smoke Exposure on Inflammatory Markers in Periphery and Brain in Young and Aged Mice

**DOI:** 10.64898/2026.02.06.703827

**Authors:** Emely A. Gazarov, Bailey McCracken, Zachary A. Krumm, Sabrina Zequeira, Barry Setlow, Jennifer L. Bizon

## Abstract

Aging is associated with chronic low-grade inflammation, which is thought to contribute to both cognitive decline and various neurodegenerative diseases. Cannabinoids are reported to reduce levels of inflammatory markers; however, these effects have not been thoroughly assessed in aged subjects. To address this gap, we evaluated effects of chronic cannabis smoke exposure on peripheral and brain inflammatory markers in young and aged mice. Young adult (4 months old) and aged (22 months old) C57Bl/6J mice were exposed to smoke from burning either cannabis (5.5 - 6.2% THC) or placebo (0% THC) cigarettes daily for 30 consecutive days. Following exposure sessions, both blood and brain tissue from the prefrontal cortex (PFC) and hippocampus (HPC) were collected and analyzed for multiple markers of inflammation.

Overall, the patterns of inflammatory markers varied across the three tissue types. Both comparisons of individual cytokines and global cytokine profiles revealed that aging caused modest increases in cytokine levels in serum and PFC, with little influence of cannabis exposure. In contrast, HPC samples had stronger age effects, with numerous cytokines elevated in aged mice compared to young. Cannabis also interacted with age in the HPC such that smoke exposure tended to increase cytokine levels in young mice but decrease them in aged mice. These findings point to general age-related increases in brain inflammatory markers in this mouse strain, but cannabis effects were largely restricted to the HPC, where smoke exposure produced age-dependent changes in cytokine profiles.

**HIGHLIGHTS:**

- Aged C57BL/6 mice have elevated cytokines levels, especially in the hippocampus
- Cannabis smoke exposure produces age-dependent changes in cytokine levels in hippocampus
- In HPC, cannabis smoke exposure attenuated age-related increases in IL-13 and Dkk1

## INTRODUCTION

Aging is associated with physiological changes in the body and brain, affecting cellular processes, vasculature, gross morphology, and ultimately cognition^1^. Consequently, aging is a major risk factor for most chronic conditions, including neurodegenerative diseases such as Alzheimer’s and Parkinson’s^2^, as well as cognitive decline not explicitly linked to neurodegeneration. These age-associated conditions commonly feature increased brain and peripheral inflammation^3^, a phenomenon linked to immunosenescence^4^. Hallmark characteristics of immunosenescence include decreased T cell production, increased levels of proinflammatory cytokines, and reduced functioning of macrophages, including microglia^1,5^. This state of chronic low-grade inflammation associated with age, or “inflammaging”, is thought to contribute to the progression of neurodegeneration and age-related cognitive deficits^6^. Studies in mice show that aged subjects exhibit elevated levels of some cytokines in the brain compared to young^7,8^, particularly in the hippocampus^7^. Similarly, human studies report that circulating levels of cytokines increase with age^9–12^.

There are increasing efforts to identify interventions that may reduce inflammatory states to mitigate risks of neurodegenerative conditions and cognitive decline. Recently, there has been growing interest in using cannabis as a therapeutic due to its anti-inflammatory and antioxidant properties^13,14^, which could be beneficial in aging. The prevalence of cannabis use among adults over 65 in the United States increased from 4.8% to 7.0% between 2021-2023^15^, and in 2021, 12.1% of adults aged 50-80 reported cannabis use in the past year^16^. Older adults often use cannabis for medical reasons, including relief from chronic pain^17^, oncological symptoms^17^, anxiety^18^, and Parkinson’s symptoms^18^. Given the rising use among this population and greater accessibility of cannabis, it is essential to understand its effects on the aging brain, particularly concerning neuroinflammation.

The two major and best studied cannabinoids found in cannabis are delta-9-tetrahydrocannabinol (THC) and cannabidiol (CBD). These cannabinoids interact with the endocannabinoid system, which includes the endogenous ligands 2-arachidonoylglycerol and anandamide that bind to two primary receptors, CB1 and CB2^19^. The brain has dense, wide-spread CB1R expression, whereas CB2Rs are predominantly found on immune cells such as microglia and in peripheral tissues^19^. Additionally, cannabinoids bind to TRPV receptors and PPARγ, which are involved in inflammatory signaling pathways^20,21^. Although microglia can release inflammatory cytokines, endocannabinoid binding can promote neural protection by shifting microglia to an alternatively activated, anti-inflammatory state^22^. Furthermore, cannabinoids can enhance microglial phagocytosis^23,24^ and migration^25^, which facilitates clearance of tissue debris and toxic protein aggregates, and promotes microglial movement to sites of injury or toxic proteins, respectively. Anti-inflammatory properties of cannabinoids were reviewed recently by Henshaw et al., in which they provided evidence from multiple studies describing CBD’s effects on reducing pro-inflammatory cytokines^13^. Notably, these authors highlighted the paucity of research on THC’s specific ability to modulate inflammatory markers.

Given the role of inflammation in aging and age-related diseases, as well as the potential for cannabinoids to attenuate inflammation, it is essential to understand how cannabinoids influence inflammation in advanced age. To date, however, research on the effects of cannabinoids on inflammatory markers has focused largely on young adult subjects and specific disease models, with few studies investigating typical aging populations^13^. Furthermore, many preclinical studies use routes of administration such as intraperitoneal injections or non-voluntary consumption with oral gavage^13^, which do not accurately model human cannabis use.

The current study aimed to address a critical gap in understanding how THC influences inflammatory signaling in both the periphery and brain in the context of aging, since older individuals represent an understudied yet increasingly relevant population of cannabis users. To investigate this, we measured inflammatory markers in serum, prefrontal cortex, and hippocampus of young and aged mice following chronic cannabis smoke inhalation.

## MATERIALS AND METHODS

### Subjects

A total of 40 C57Bl/6J mice (half female) were obtained from Jackson Laboratories. At the onset of smoke exposure, the mice were either 4 months (n = 20) or 22 months (n = 20) old, resulting in 10 mice per sex per age group. The mice were housed single sex, 5 per cage, under a 12-hr light/dark cycle (lights on at 0700) with the vivarium maintained at 25°C. All experiments were conducted during the light phase. Water and food (2918 Teklad global 18% protein diet) were provided *ad libitum*. All animal procedures were approved and performed in accordance with the University of Florida Institutional Animal Care and Use Committee (protocol number IACUC202200000499) and followed National Institutes of Health guidelines.

### Experimental design

For each sex and age combination, 5 mice were randomly assigned to undergo daily exposure to smoke generated from sequentially burning either 3 placebo or 3 cannabis cigarettes (approximately 700 mg per cigarette; NIDA Drug Supply Program), for a total of 30 days. Placebo cigarettes contained ≤ 0.1% w/w THC or any other cannabinoids, while cannabis cigarettes contained 5.5-6.2% w/w THC (approximately 40 mg THC per cigarette) and minimal levels of other cannabinoids, including CBD, CBN, and CBC (≤ 0.5% w/w). Each smoke exposure session lasted approximately 40 minutes (the time required to burn the 3 cigarettes in their entirety).

Smoke exposure sessions were conducted using a TE-10 automated cigarette smoking machine (Teague Enterprises, Davis, CA USA). The procedures and validation of reliable THC and THC metabolite levels in plasma and brain resulting from this exposure regimen are detailed in our previous work^26–28^. Briefly, during exposure sessions, mice remained in their home cages, which were placed in the exposure chamber of the machine. Cannabis cigarettes were lit and puffed (35 cm^3^ puff volume, 1 puff per min, 2 s per puff) in the ignition chamber, from which the smoke produced was pumped into the exposure chamber and then exhausted to the exterior of the building.

On day 31, mice were euthanized via rapid decapitation. Trunk blood was collected using Microvette^®^ CB 300 Serum and Clotting Activator tubes (Sarstedt Inc, Newton, NC USA). Blood was allowed to coagulate for 20 min at room temperature, then temporarily stored on ice. Blood was then centrifuged for 10 min at 4500 rpm (1902 x g) at 4°C to separate serum. Serum was pipetted and transferred to polypropylene microcentrifuge tubes and stored at −80°C until protein quantification. Brains were harvested and flash frozen in 2-methylbutane chilled in a dry ice/ethanol bath for approximately 30 s. The frozen brains were temporarily stored on dry ice during harvesting, and then stored at −80°C until sectioning.

### Sample preparation and protein quantification

Frozen serum samples were analyzed by a commercial vendor (Raybiotech, Peachtree Corners, GA, USA) using a custom quantitative sandwich-based antibody array to test for 40 cytokines (**Table 1**).

**Table 1.**
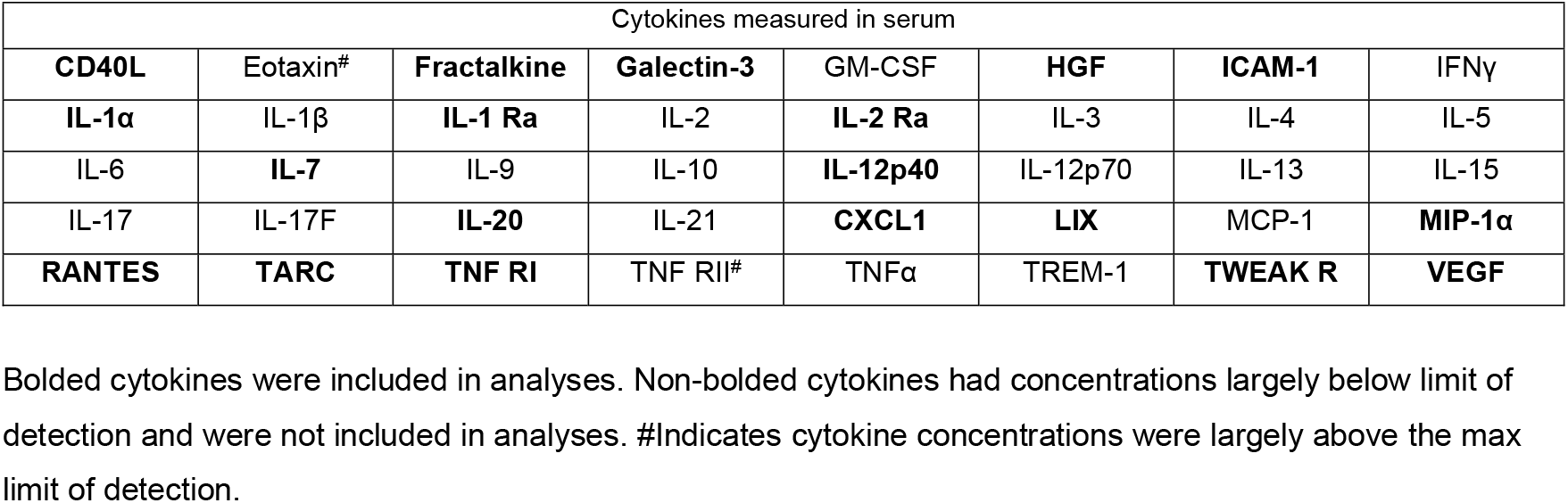
Serum Cytokines.

Prior to sectioning, brains were transferred to a −20°C freezer overnight. The brains were then cryosliced, and tissue from the prefrontal cortex (PFC) [including prelimbic, infralimbic, anterior cingulate, and orbitofrontal cortices] and hippocampus (HPC) [including dentate gyrus and CA1, CA2, and CA3 subregions] was collected using 1 mm biopsy punches (Integra Miltex, York, PA USA). Tissue from each region was homogenized in lysis buffer (50 mM HEPES, 1mM EDTA, 1 mM EGTA, pH 7.4) containing a protease and phosphatase inhibitor (Pierce Protease and Phosphatase Inhibitor Mini Tablets) and stored at −80°C until protein isolation.

Brain tissue homogenates were centrifuged for 5 min at 10,000 x g at 4°C, and the supernatants were collected into clean polypropylene tubes. The protein concentrations of all PFC and HPC brain lysates were determined using the Pierce 660 nm protein assay^29^ by generating an 8-point BSA standard curve, with each sample run in triplicate. To ensure reliability, R^2^ values for standard curves were ≥0.996 and no sample had a coefficient of variation (CV) greater than 10%. Once concentrations were determined, samples were diluted to standardize each to 90 μL of 1.5 mg/mL protein. Frozen PFC and HPC lysates were analyzed by a commercial vendor (Raybiotech, Peachtree Corners, GA, USA) using a quantitative sandwich-based antibody array (QAM-CAA-2000-1) to determine levels of 120 inflammatory markers (**Table 2)**.

**Table 2.**
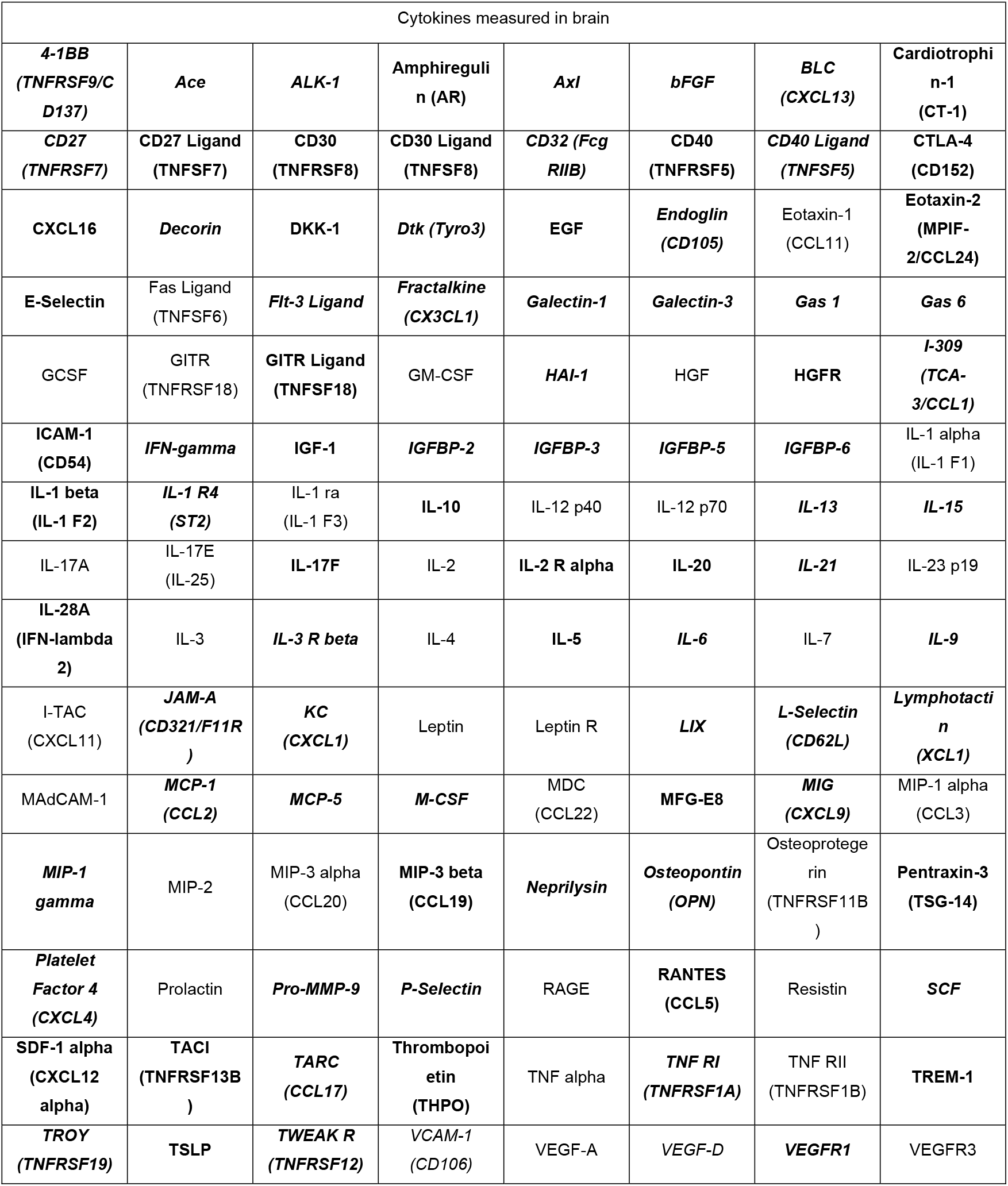
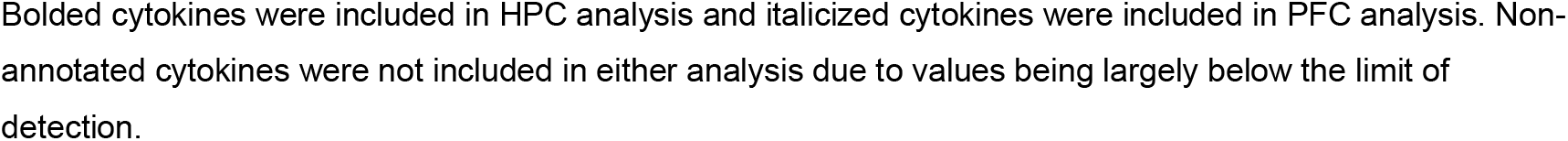
PFC and HPC Cytokines.

| Cytokines measured in brain |  |  |  |  |  |  |  |
| --- | --- | --- | --- | --- | --- | --- | --- |
| **4-1BB**<br>(*TNFRSF9/C*<br>*D137*) | *Ace* | *ALK-1* | **Amphiregulin**<br>(AR) | *Axl* | *bFGF* | **BLC**<br>(*CXCL13*) | **Cardiotrophin-1**<br>(CT-1) |
| **CD27**<br>(*TNFRSF7*) | **CD27 Ligand**<br>(*TNFSF7*) | **CD30**<br>(*TNFRSF8*) | **CD30 Ligand**<br>(*TNFSF8*) | **CD32** (*FcgRIIB*) | **CD40**<br>(*TNFRSF5*) | **CD40 Ligand**<br>(*TNFSF5*) | **CTLA-4**<br>(CD152) |
| **CXCL16** | *Decorin* | **DKK-1** | *Dtk* (*Tyro3*) | **EGF** | *Endoglin*<br>(*CD105*) | Eotaxin-1<br>(*CCL11*) | **Eotaxin-2**<br>(*MIPF-2/CCL24*) |
| **E-Selectin** | Fas Ligand<br>(*TNFSF6*) | *Flt-3 Ligand* | *Fractalkine*<br>(*CX3CL1*) | *Galectin-1* | *Galectin-3* | **Gas 1** | **Gas 6** |
| GCSF | GITR<br>(*TNFRSF18*) | **GITR Ligand**<br>(*TNFSF18*) | GM-CSF | *HAI-1* | HGF | **HGFR** | *I-309*<br>(*TCA-3/CCL1*) |
| **ICAM-1**<br>(CD54) | *IFN-gamma* | **IGF-1** | *IGFBP-2* | *IGFBP-3* | *IGFBP-5* | *IGFBP-6* | IL-1 alpha<br>(IL-1 F1) |
| **IL-1 beta**<br>(IL-1 F2) | **IL-1 R4**<br>(*ST2*) | IL-1 ra<br>(IL-1 F3) | **IL-10** | IL-12 p40 | IL-12 p70 | **IL-13** | **IL-15** |
| IL-17A | IL-17E<br>(IL-25) | **IL-17F** | IL-2 | **IL-2 R alpha** | **IL-20** | **IL-21** | IL-23 p19 |
| **IL-28A**<br>(IFN-lambda 2) | IL-3 | *IL-3 R beta* | IL-4 | **IL-5** | **IL-6** | IL-7 | **IL-9** |
| I-TAC<br>(*CXCL11*) | *JAM-A*<br>(*CD321/F11R*) | **KC**<br>(*CXCL1*) | Leptin | Leptin R | **LIX** | **L-Selectin**<br>(*CD62L*) | **Lymphotactin**<br>(*XCL1*) |
| MAdCAM-1 | **MCP-1**<br>(*CCL2*) | **MCP-5** | **M-CSF** | MDC<br>(*CCL22*) | **MFG-E8** | **MIG**<br>(*CXCL9*) | MIP-1 alpha<br>(*CCL3*) |
| **MIP-1 gamma** | MIP-2 | MIP-3 alpha<br>(*CCL20*) | **MIP-3 beta**<br>(*CCL19*) | *Neprilysin* | **Osteopontin**<br>(*OPN*) | Osteoprotegerin<br>(*TNFRSF11B*) | **Pentraxin-3**<br>(*TSG-14*) |
| **Platelet Factor 4**<br>(*CXCL4*) | Prolactin | **Pro-MMP-9** | **P-Selectin** | RAGE | **RANTES**<br>(*CCL5*) | Resistin | **SCF** |
| **SDF-1 alpha**<br>(*CXCL12* alpha) | **TACI**<br>(*TNFRSF13B*) | **TARC**<br>(*CCL17*) | **Thrombopoietin**<br>(*THPO*) | TNF alpha | **TNF RI**<br>(*TNFRSF1A*) | TNF RII<br>(*TNFRSF1B*) | **TREM-1** |
| **TROY**<br>(*TNFRSF19*) | **TSLP** | **TWEAK R**<br>(*TNFRSF12*) | *VCAM-1*<br>(*CD106*) | VEGF-A | *VEGF-D* | **VEGFR1** | VEGFR3 |

### Statistical analysis

In order to have group sizes that were sufficient for meaningful analysis, male and female mice were combined within each Age x Drug group, for a total of n = 10/group. For each sample type (serum, PFC, and HPC), only cytokines with seven or more detectable values in at least one of the four Age x Drug groups were included in the analysis. For those cytokines that met this inclusion criterion, values below the limit of detection (LOD) in individual mice were imputed as ½ LOD, and values above the quantifiable range were imputed as the assay’s maximum measurable value. Cytokine concentrations were Log_2_-transformed to normalize distribution and satisfy model assumptions. Technical outliers, defined as values exceeding 4 standard deviations (SD) from the global mean of that cytokine, were winsorized to the ±4 SD threshold. Multivariate outliers were identified using Principal Component Analysis (PCA) based on Euclidean distance to group centroids; subjects exceeding the interquartile range threshold (Q3 + 2 x IQR or Q1 – 2 x IQR) were entirely excluded from all analyses within that sample type, including the individual and global analysis. These exclusions consisted of two mice from the serum dataset (2 Aged Cannabis mice) and one mouse from the HPC dataset (Young Placebo). No outliers were identified for PFC samples.

The goal of the analysis sequence was to assess both individual cytokine changes and global cytokine profiles. To determine the effects of age and cannabis smoke exposure on individual cytokines, we used two-way analysis of variance (ANOVA) with Age (Young, Aged) and Drug (Placebo, Cannabis) as fixed factors. ANOVAs were performed using ordinary least squares with Type III sums of squares and sum contrasts. To control for multiple comparisons, a 5% false discovery rate correction (FDR, q = 0.05) was applied independently to each factor across all cytokines. Significant Age x Drug interactions were followed by post-hoc pairwise comparisons using Welch’s t-tests with Bonferroni correction. Heatmaps of Z-scored cytokine values were generated using hierarchical clustering on rows to visualize co-regulated clusters across Age x Drug groups.

To characterize global cytokine profiles, we applied PCA to Z-scored data using the overall mean and SD for each cytokine across all mice within each sample type. The effects of Age, Drug, and their interaction on PC1 and PC2 coordinates were evaluated using two-way ANOVA (Type III Sum of Squares). To identify the primary drivers of these global cytokine profiles, we examined the top five loading magnitudes (absolute values) for PC1 and PC2. For top contributors not previously identified in the FDR-corrected individual analyses, exploratory two-way ANOVAs were conducted to identify underlying trends. Data processing and statistical analysis were conducted in Python (Jupyter Notebook/Google Colaboratory) and GraphPad Prism (Version 10).

## RESULTS

### Individual Cytokine Analyses

To examine the effects of age and cannabis smoke exposure on individual cytokine levels, we conducted ANOVA analyses across serum, PFC, and HPC samples. While serum and PFC cytokines showed age-related changes, HPC exhibited both age-related increases in inflammatory markers and several alterations associated with cannabis smoke exposure.

For serum, data from 19/40 cytokines were analyzed, with the remaining cytokines falling below the limit of detection or above the maximum (**Table 1**). For PFC and HPC, data were analyzed for 55/120 and 85/120 cytokines, respectively (**Table 2**). To visualize patterns of cytokine changes, heat maps for serum (**Fig. 1A**), PFC (**Fig. 1B**), and HPC (**Fig. 1C**) were generated from Log_2_ transformed cytokine levels, with each cytokine z-scored across all mice. Two-way ANOVA with FDR correction were performed on each Log_2_ transformed cytokine to assess the effects of age and cannabis smoke exposure on individual cytokines (adjusted p-values are reported). For serum samples, there was a significant main effect of Age on IL-12p40 (F(1,34) = 14.4, p = 0.011) and RANTES (F(1,34) = 12.7, p = 0.011) levels (**Table 3**), such that they were greater in aged mice compared to young. In PFC, there were significant main effects of Age on Galectin-3 (F(1,36) = 32.3, p = 0.0001), PF4 (F(1,36) = 22.4, p = 0.0009, KC (F(1,36) = 16.0, p = 0.005), OPN (F(1,36) = 13.5, p = 0.01), P-selectin (F(1,36) = 13.1, p = 0.01), and MIP-1γ (F(1,36) = 11.6, p = 0.015), such that aged mice had higher levels of these cytokines compared to young (**Table 3**). In HPC, there was a significant main effect of Age on P-Selectin (F(1,35) = 37.3, p < 0.0001), PF4 (F(1,35) = 34.2, p < 0.0001), TNF R1 (F(1,35) = 16.9, p = 0.006), and bFGF (F(1,35) = 13.5, p = 0.02) with all cytokine levels higher in aged mice compared to young (**Table 3**).

**Table 3.**
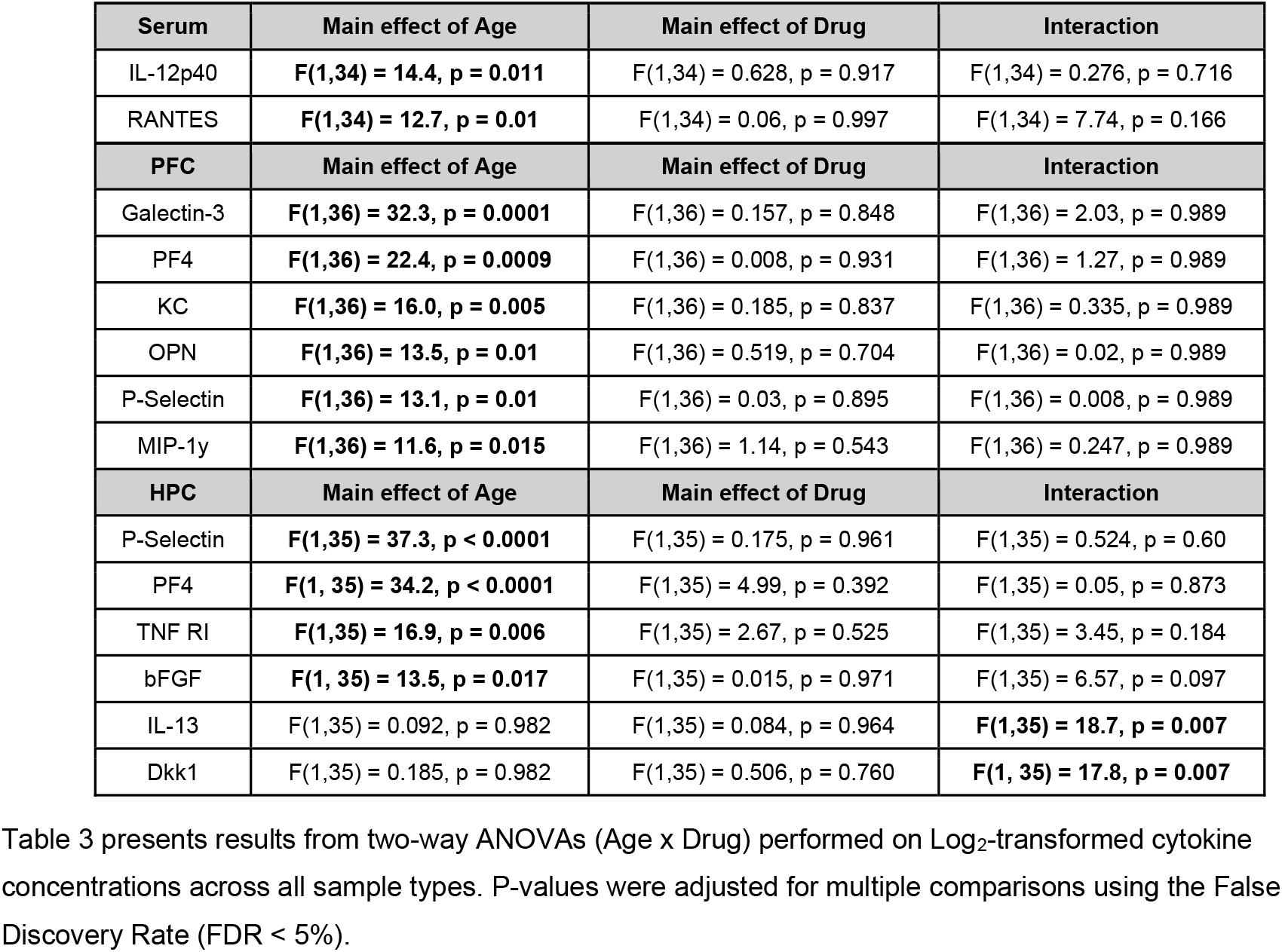
Individual Cytokine Analysis.

**Figure 1.**
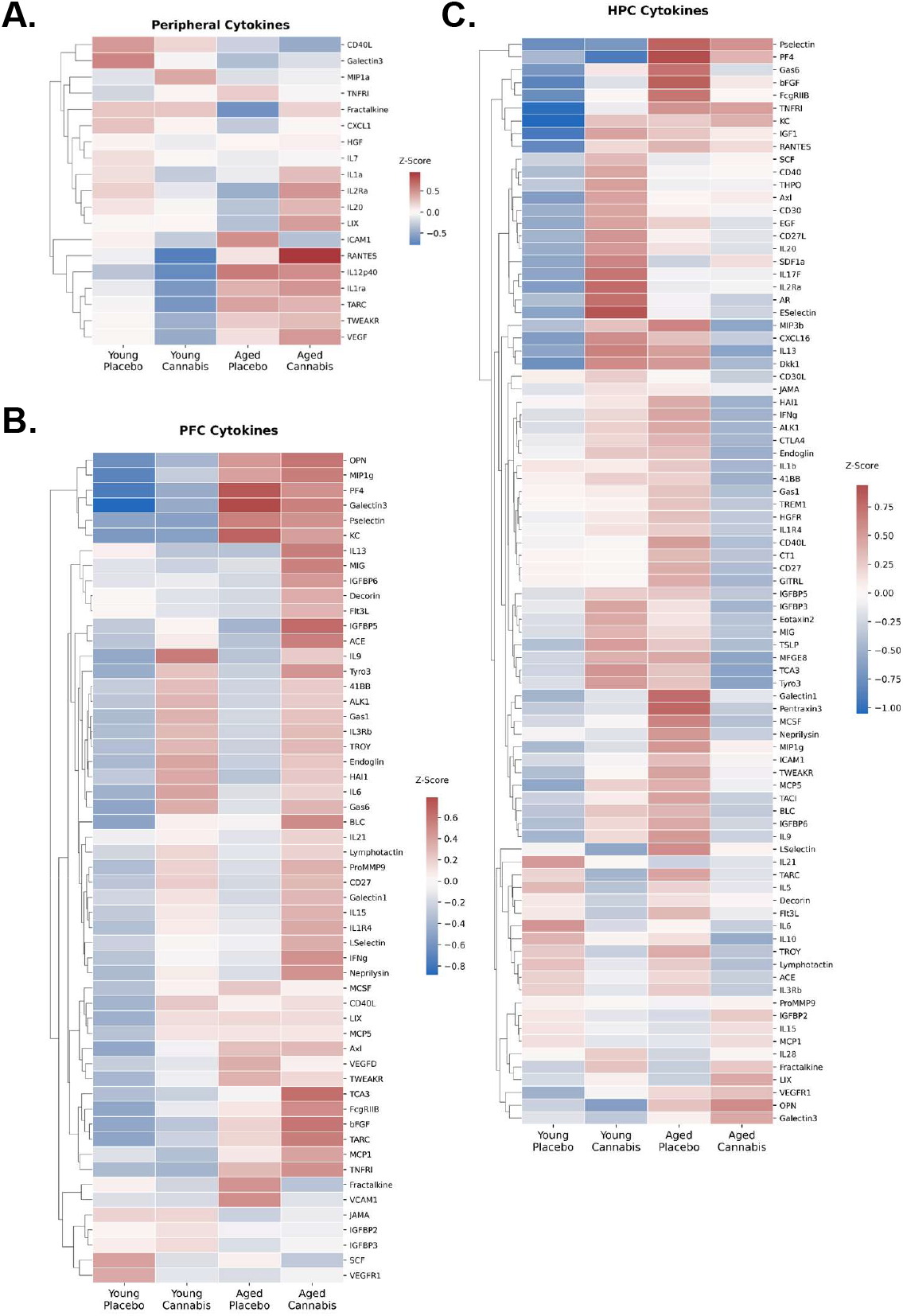
Cytokine Heatmaps. For each sample type, z-scores were calculated for each Log_2_-transformed cytokine using the overall mean and SD across all mice. Each cell shows the mean z-score for a given Age x Drug group, with the color bar indicating z-score magnitude and direction. Hierarchical clustering was performed to order cytokines and experimental conditions by similarity. Heatmaps display averaged z-scores for serum **(A)**, Prefrontal Cortex **(B)**, and Hippocampus samples **(C)**.

In addition to these main effects of Age on HPC cytokines, there were significant Age x Drug interactions for the following cytokines in HPC: IL-13 (F(1,35) = 18.7, p = 0.007) and Dkk1 (F(1,35) = 17.8, p = 0.007 (**Table 3**). Post hoc multiple comparisons with Bonferroni correction revealed that aged placebo mice had significantly higher levels of IL-13 (p = 0.025) and Dkk1 (p = 0.008) compared to young placebo mice. In addition, cannabis exposure had age-specific effects such that aged cannabis mice had significantly lower levels of IL-13 (p = 0.02) and Dkk1 (p = 0.0498) compared to aged placebo mice, suggesting that it reversed the age-related increases in these cytokines. Moreover, young cannabis mice had significantly higher levels of IL-13 (p = 0.008) and Dkk1 (p = 0.005) compared to young placebo mice, suggesting that cannabis smoke exposure has opposite effects in young and aged mice in HPC. See Supplementary Figure 1 for bar graphs corresponding to the data in Table 3.

### Principal Component Analyses of Cytokine Profiles

While the initial ANOVA analyses examined cytokines individually, this approach may overlook coordinated changes across multiple cytokines. To capture these broader changes in cytokine levels, we used PCA to assess how aging and cannabis exposure shape cytokine profiles in each sample type. PCA was performed on Z-scored cytokines for each sample type (serum, PFC, and HPC). We visualized individual samples and group centroids in PCA space to examine clustering patterns. The five cytokines with the highest absolute loading values on the first two principal components were identified as potential drivers of the variance of each axis. See Supplementary Figure 2 for bar graphs displaying loading values for all cytokines within each sample type. To further explore these top contributors, we conducted two-way ANOVAs (Age x Drug) with post-hoc multiple comparisons with Bonferroni corrections (adjusted p values are reported) on cytokines that had not been previously identified in the individual analyses. Additionally, we determined the effects of Age, Drug, and their interaction on principal components (PC) axes.

For serum samples, PCA (**Fig. 2A**) revealed that PC1 explains 24.4% of the total variance, with VEGF, IL2Ra, TWEAKR, TARC, and IL-12p40 as the top five cytokine contributors (**Fig. 2B**); however, PC1 was not significantly affected by Age, Drug, or interaction effects. As previously described, IL-12p40 levels were higher in aged mice compared to young, and exploratory two-way ANOVAs showed a significant main effect of Age on TARC (F(1,34) = 5.46, p = 0.026), such that levels of these cytokines were higher in aged mice compared to young. PC2 explained 12.6% of the total variance and was also not significantly affected by Age, Drug, or interaction effects. The top 5 cytokines contributing to PC2 were IL-1α, IL20, IL-1Ra, MIP1α, and TWEAKR (**Fig. 2C**). There was a main effect of age on IL-1Ra (F(1,34) = 6.29, p = 0.017) such that levels of these cytokines were higher in aged mice compared to young. Although these cytokines were not all statistically significant in individual ANOVA analysis of age effects, they emerged as the top contributors in the PCA. Given that the PCA captures shared patterns of variation across cytokines, this suggests that these cytokines may be a part of a global, age-associated immune signature not evident in univariate analyses alone.

**Figure 2.**
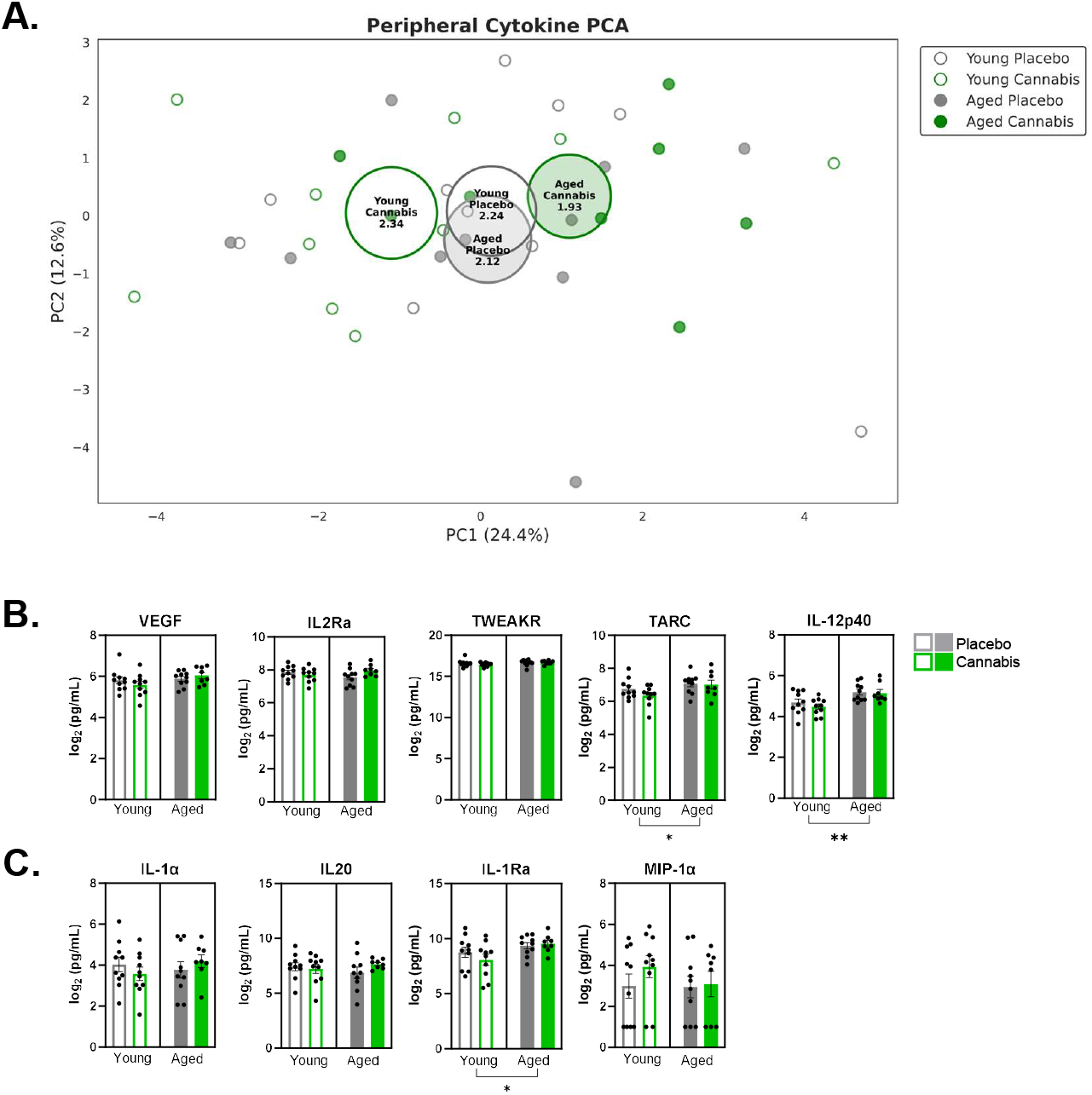
Principal component analysis of serum cytokine profiles. PCA plot showing individual mouse values and group centroids by Age and Drug group **(A)**. Log_2_-transformed cytokine concentrations of the top five PC1 contributors: VEGF, IL2Ra, TWEAKR, TARC, and IL-12p40, plotted by Age x Drug group **(B)**. Log_2_-transformed cytokine concentrations of the top five PC2 contributors: IL-1α, IL20, IL-1Ra, MIP-1α, and TWEAKR, plotted by Age x Drug group **(C)**. While PC1 and PC2 were not significantly affected by Age or Drug, several individual cytokines showed significant main effects, suggesting more targeted rather than global effects of cannabis smoke on cytokine expression.

For PFC samples, PCA (**Fig. 3A**) revealed that PC1 explained 32.9% of the total variation, with the top five contributing cytokines including CD40L, Gas1, IL1R4, ALK1, and Endoglin (**Fig. 3B**). PC2 explained 14.4% of the total variance and the top contributors were TCA3, PF4, MIP1γ, KC, and MIG (**Fig. 3C**). While principal component coordinates for PC1 were not significantly affected by Age, Drug, or their interaction, there was a significant effect of Age on PC2 coordinates (F(1,34) = 8.33, p = 0.007). Additional analysis of the top five cytokines only revealed main effects of Age on cytokines previously identified in the individual analysis (PF4, MIP1γ, and KC).

**Figure 3.**
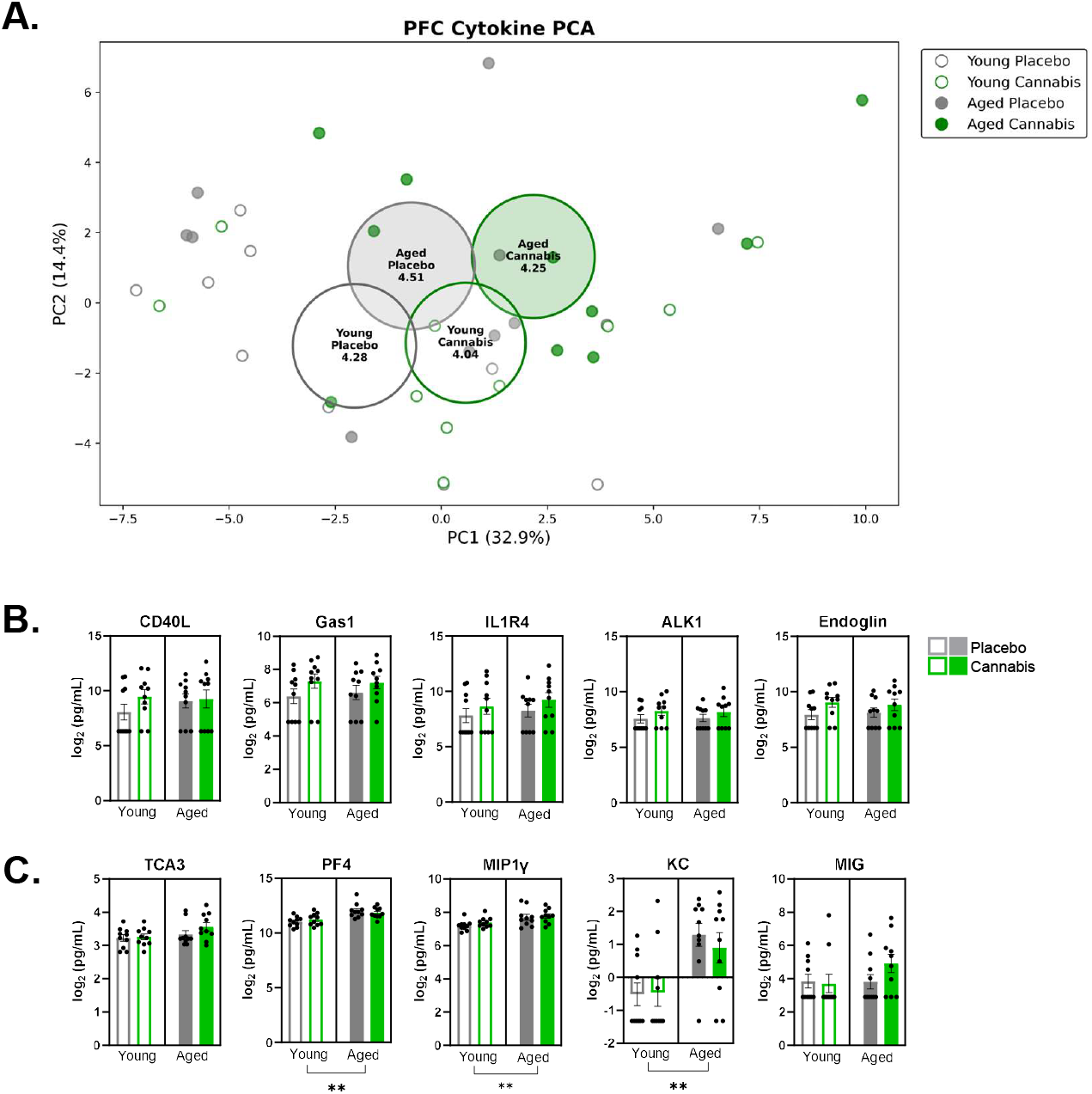
Principal component analysis of prefrontal cortex cytokine profiles. PCA plot showing individual mouse values and group centroids by Age and Drug group **(A)** Log_2_-transformed cytokine concentrations of the top five PC1 contributors: CD40L, Gas1, IL1R4, ALK1, Endoglin, plotted by Age x Drug group **(B)**. Log_2_-transformed cytokine concentrations of the top five PC2 contributors: TCA3, PF4, MIP-1γ, KC, and MIG, plotted by Age x Drug group **(C)**. PC2 was significantly affected by Age (p = 0.007), which is consistent with the age-related differences observed in the individual analysis of its top contributing cytokines.

In contrast to the results of the PCAs in serum and PFC samples, PCA of HPC cytokine profiles revealed a significant Age x Drug interaction on PC1 coordinates (F(1,35) = 8.96, p = 0.005), with this axis accounting for 25.5% of the total variance (**Fig. 4A**). Additionally, PC2, which explained 18.4% of the total variance, was significantly affected by Drug (F(1,35) = 4.99, p = 0.03). The influence of the Age x Drug interaction on PC1 was further supported by exploratory two-way ANOVAs conducted on the top contributing cytokines, which included ALK1, IL-1R4, TACI, Gas6, and CD40L (**Fig. 4B**). There were significant Age x Drug interactions observed for ALK1 (F(1,35) = 4.65, p = 0.038) and Gas6 (F(1,35) = 9.56, p = 0.004) levels. Post hoc comparisons with Bonferroni correction showed that aged placebo mice had higher levels of Gas6 (p = 0.004) compared to young placebo mice. Significant interaction effects were also observed for the top five cytokines contributing to PC2, which included IL-2Ra, Amphiregulin, IL-3RB, EGF, and IL20 (**Fig. 4C**). Significant Age x Drug interactions were seen for IL-2Ra (F(1,35) = 5.66, p = 0.023), Amphiregulin (F(1,35) = 5.66, p = 0.023), EGF (F(1,35) = 5.44, p = 0.026), and IL20 (F(1,35) = 5.05, p = 0.031). Post hoc tests showed that young cannabis mice had significantly higher levels of IL-2Ra (p = 0.004), and Amphiregulin (p = 0.026) compared to young placebo mice. In combination (and consistent with the results of the comparisons of individual cytokines), these results suggest that cannabis smoke exposure has age-specific effects on HPC cytokine profiles, such that cannabis tends to decrease levels of HPC cytokines in aged mice but increases them in young mice.

**Figure 4.**
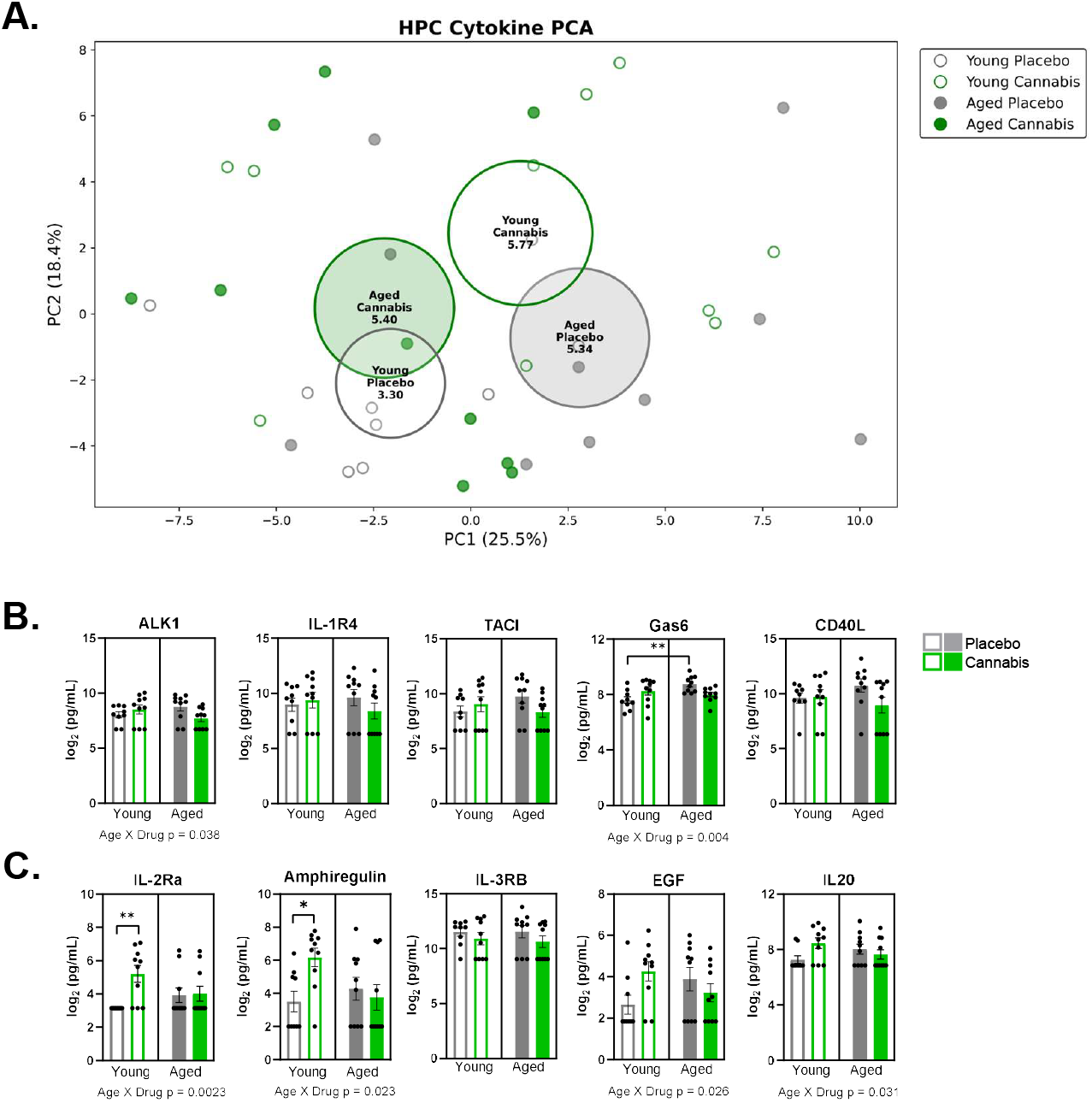
Principal component analysis of hippocampal cytokine profiles. PCA plot showing individual mouse values and group centroids by Age and Drug group **(A)**. Log_2_-transformed cytokine concentrations of the top five PC1 contributors: ALK1, IL1R4, TACI, Gas6, and CD40L, plotted by Age x Drug group **(B)**. Log_2_-transformed cytokine concentrations of the top five PC2 contributors: IL2Ra, Amphiregulin, IL3Rb, EGF, and IL20, plotted by Age x Drug group **(C)**. PC1 was significantly influenced by the Age x Drug interaction (p = 0.005), and there was a significant effect of Drug on PC2 (p = 0.03). Exploratory analyses of top contributing cytokines found that cannabis smoke exposure decreased levels of multiple cytokines in aged mice and increased them in young mice, supporting these global interaction effects.

## DISCUSSION

This study compared both peripheral and brain inflammatory profiles in young and aged mice, and determined how chronic cannabis smoke exposure influenced these profiles. With growing interest in the therapeutic potential of cannabis and increased use among older adults, understanding the interaction between aging and cannabis use is critical. While research on the anti-inflammatory properties of cannabis has largely focused on CBD, the immunological effects of THC remain poorly understood. To address this, we used a validated smoke exposure protocol using THC-dominant cannabis to ensure effective delivery of THC into both the brain and systemic circulation of young and aged mice^26,28^.

### Peripheral cytokines

Human aging is associated with increased levels of circulating cytokines, including IL-12p70^9^, TNFα^9,10^, IL-6^9–12^, IL-10^9,10^, TGF-α^30^, and IL-1Ra^31^. Similarly, age-related changes in circulating cytokines are observed in rodent models, such that aged mice have elevated serum levels of Eotaxin, IL-9, and TARC compared to young mice^32^. Although the specific cytokines that vary with age can differ between studies, likely in part due to differences in blood collection methods, assay sensitivity, and technical variability^33^, there is consistent evidence in rodents and humans that aging is accompanied by increased systemic inflammation.

Consistent with the literature, our current study revealed that aged mice had significantly higher serum levels of IL-12p40 and RANTES compared to young mice. Notably IL-12p40 emerged as a top contributor in our global cytokine analysis, suggesting its potential utility as a peripheral biomarker for mouse aging. Additionally, exploratory analyses showed significant age-related increases in TARC and IL-1Ra levels, which is in agreement with existing findings^31,32^. Although there were no significant main or interaction effects of cannabis smoke exposure on serum cytokines, the age-related increase in RANTES is of particular interest. RANTES is a pro-inflammatory cytokine known to mediate leukocyte migration^34^. While prior *in vitro* studies indicate that THC can inhibit macrophage chemotactic responses to RANTES^35^, our chronic smoke exposure protocol did not significantly alter circulating RANTES levels. This suggests that THC’s immunomodulatory effects on RANTES may be tissue-specific or dose-dependent, and further research should investigate these mechanisms.

### Central Cytokines

Inflammaging is known to play a role in age-related cognitive decline. Given the contributions of the PFC and HPC in executive and mnemonic functions, respectively, we investigated region-specific inflammatory profiles in young and aged mice. Prior studies in mice have reported increased cytokine levels in these brain regions with aging. Porcher et al. found that IL-1α, IL-1β, and IL-17 were upregulated in the hippocampus of aged mice^7^, while Cyr & de Rivero Vaccari reported increased levels of IL-1α, IL-8, IL-17a, IL-7, LT-α, and VEGF-A in the cortex of aged mice compared to young^8^.

#### Prefrontal Cortex

In the current study, we found that Galectin-3, PF4, KC, OPN, P-Selectin, and MIP-1γ levels were significantly elevated in PFC of aged compared to young mice. Additionally, PF4, KC, and MIP-1γ, were identified in the global analysis, suggesting that they are likely drivers of age-related inflammation in mouse PFC. A number of these markers, including Galectin-3 and KC, have also been shown to play a role in the pathogenesis of neurodegenerative conditions such as Alzheimer diseases^36–38^, by modulating microglial activation and inducing tau cleavage, respectively. Interestingly, circulating PF4 is reported to decline with age and systemic administration of PF4 in aged mice has been shown to improve cognitive function^39^; however, we found that brain PF4 levels were elevated in aged mice. The discrepancy between peripheral and central PF4 levels highlights the need for further research on this cytokine in the context of aging. Cannabis exposure did not significantly alter global cytokine levels in the PFC; however, trends (0.05 < p < 0.1) toward increased Gas1, ALK1, and Endoglin in cannabis exposed mice point to possible modulatory effects. This pattern was reinforced with a trending main effect of Drug on PC1 (p = 0.095). Future studies with larger sample sizes should explore these markers to determine their potential roles in cannabis-related changes in inflammation.

#### Hippocampus

Analyses of hippocampal cytokines revealed effects of both age and cannabis smoke, highlighted by a significant Age x Drug interaction in the global cytokine analysis. Regarding the effects of age alone, the findings revealed higher levels of several cytokines in the aged HPC, including P-Selectin, PF4, TNF RI, and bFGF. Additionally, both the individual and global analyses identified significant Age x Drug interactions for cytokines including IL-13, Dkk1, and ALK1.

These changes hold relevance for aging since Dkk1, a Wnt pathway antagonist, is known to increase in the HPC with age and contributes to synapse loss and cognitive decline^40–42^. Interestingly, a recent genome-wide association study using human whole blood samples reported that long-term cannabis use in middle aged adults is associated with alterations in expression of genes linked to WNT signaling pathways^43^, supporting the translational relevance of our findings. Similarly, ALK1 is a component of the TGF-β complex, which plays a role in pathological aging and neuroinflammation^44^. The interaction effect suggests that cannabis may modulate components of this signaling pathway to exert anti-inflammatory effects in aging. This aligns with prior evidence that THC reduced connective tissue growth factor (CTGF), a TGF-β modulator, in aged mice^45^, and emphasizes further need to study the effects of THC on this signaling pathway.

Post-hoc tests showed that cannabis exposure reduced IL-13 and Dkk1 in aged mice, whereas young mice exposed to cannabis exhibited increased levels of IL-13, Dkk1, IL-2Ra and Amphiregulin. This bidirectional response helps explain the differential behavioral effects of cannabis on young and aged subjects, which has previously been demonstrated in our lab^28^ and others^46,47^. Recent work has shown that IL-13 modulates VTA dopamine neuron firing^48^, which offers one possible mechanism by which these neurobiological changes in immune signaling could lead to age-dependent behavioral outcomes. Consistent with our findings, HPC-specific effects of THC in aged subjects have previously been demonstrated by Bilkei-Gorzo et al. (2017). In that study, aged mice treated with chronic low dose THC showed hippocampal gene expression patterns that closely resemble those of young control mice^45^. Additionally, young mice treated with THC had impaired hippocampal-dependent cognitive performance, whereas aged mice showed improvements. The neurobiological changes in HPC immune signaling may help explain why cannabis has differential effects on behavior in young and aged subjects. Future studies should investigate the causal role of these specific cytokines, such as IL-13 and Dkk1, in mediating the age-dependent effects of cannabis on cognition.

### Limitations and Conclusions

The present study has several limitations that should be acknowledged. First, blood samples were collected only at the experimental endpoint, which prevented assessment of baseline cytokine levels prior to cannabis or placebo smoke exposure. Although this limits our ability to track within-subject changes over time, the need for brain tissue collection made a single terminal timepoint the most feasible approach. Second, the study was not adequately powered to assess sex differences. To ensure sufficient statistical power for age and drug treatment comparisons, male and female data were combined in all analyses. Prior work has demonstrated sex-specific patterns of neuroinflammation in aged mice, particularly in the HPC, in that cytokine increases with age are more pronounced in females^7^. Future research should assess sex differences in cannabis-induced immune modulation. Additionally, because smoke inhalation itself is known to increase proinflammatory cytokines in lung tissue and the frontal cortex^49,50^, we utilized a placebo smoke control to isolate the specific effects of THC from the inflammatory impact of combustion products. Future studies should investigate the effects of THC on inflammatory markers using alternate routes of administration, such as edible consumption.

The primary goal of this study was to identify candidate cytokines that may mediate age-dependent effects of chronic cannabis smoke exposure. Our results revealed that while multiple cytokines were elevated in aged mice relative to young mice across serum, PFC, and HPC, the effects of cannabis were largely specific to the HPC. In particular, the significant Age x Drug interaction in global cytokine profiles indicated that cytokine expression increased with age in placebo control mice, and that cannabis smoke exposure reduced cytokine levels in aged mice, while increasing them in young mice. These bidirectional effects may help explain the differential impact of cannabis on cognition observed in younger versus older subjects^28,45,51^. These findings provide promising molecular targets for future investigation to elucidate the causal role of specific inflammatory markers in age- and cannabis-related changes in cognitive function.

## ACKNOWLEDGEMENTS

We thank Brandon Hellbusch for assistance with sample preparation, and the Drug Supply Program at the National Institute on Drug Abuse for providing cannabis and placebo cigarettes.

## AUTHORSHIP CONTRIBUTIONS

Emely Gazarov: Investigation, Data Analysis, Writing – Original Draft Preparation; Bailey McCracken: Investigation, Writing – Original Draft Preparation; Zak Krumm – Data Analysis; Sabrina Zequeira: Investigation; Barry Setlow: Conceptualization, Resources, Supervision, Methodology, Writing – Review and Editing; Jennifer L. Bizon: Conceptualization, Resources, Supervision, Methodology, Writing – Review and Editing

