## Supplementary figures and images for "Effects of Chronic Cannabis Smoke Exposure on Inflammatory Markers in Periphery and Brain in Young and Aged Mice"

### Supplemental Figures

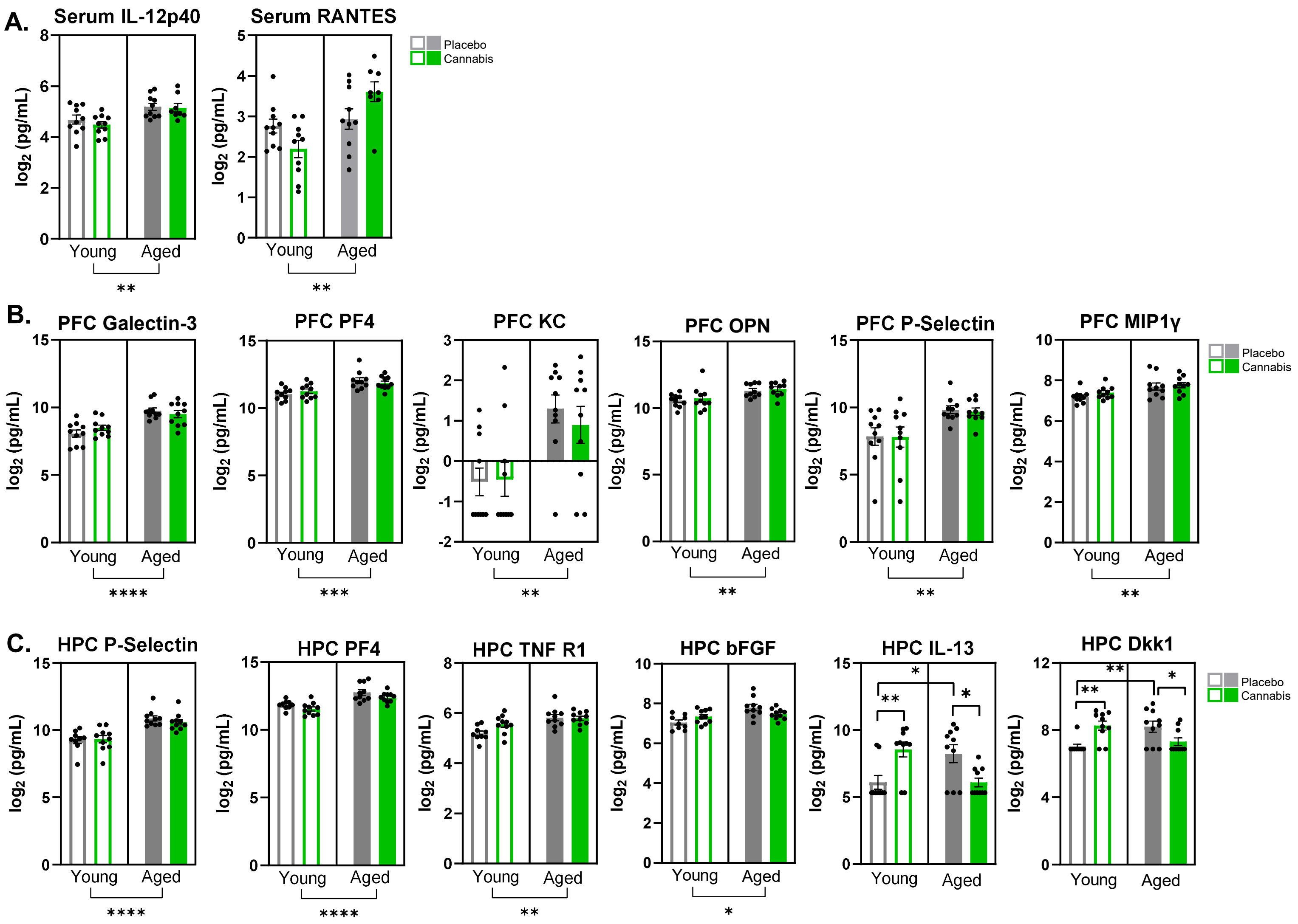

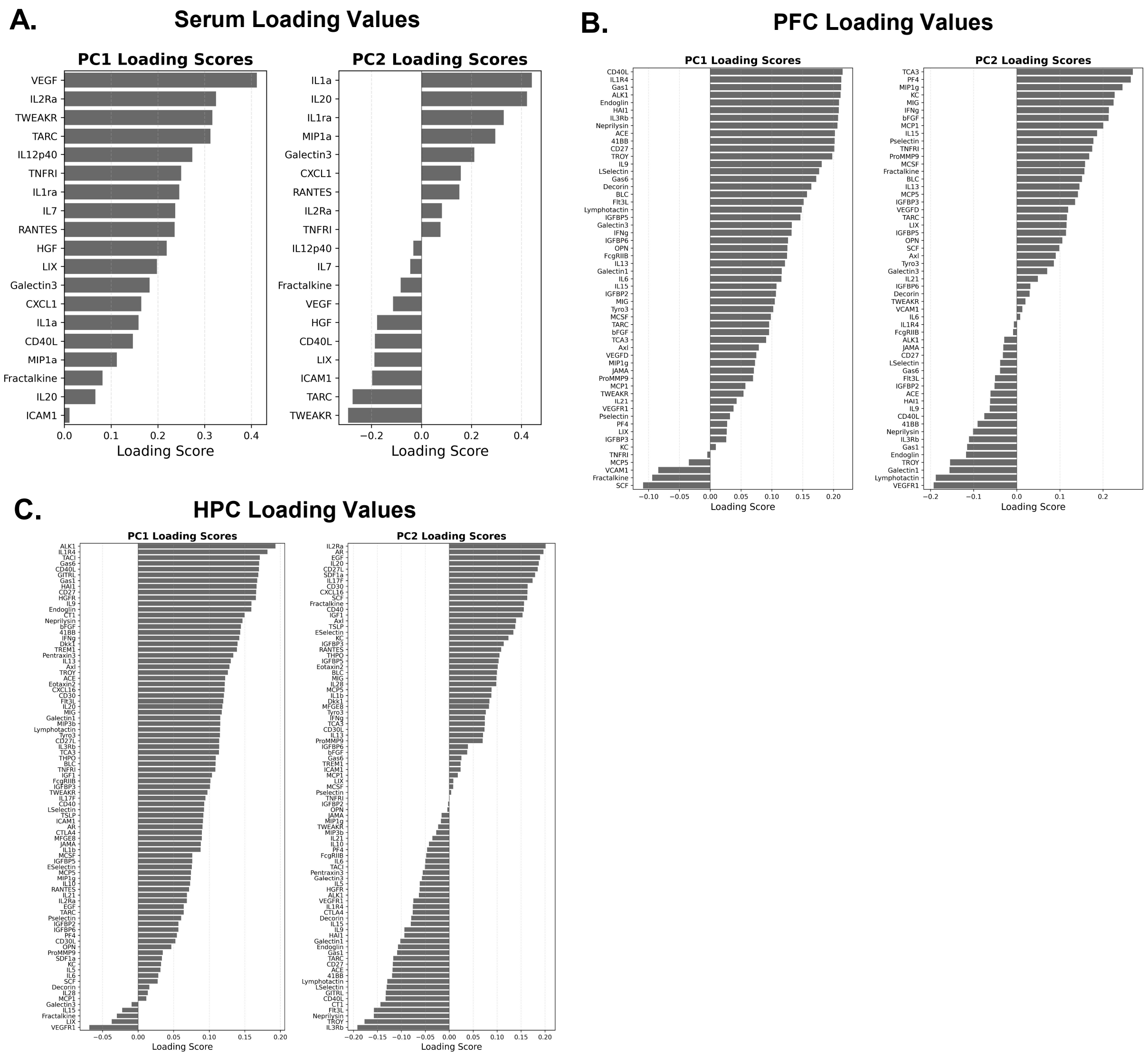
